# Hitting the right pitch: Cortical tracking of speech fundamental frequency in auditory and somatomotor regions

**DOI:** 10.1101/2025.03.18.644060

**Authors:** Yorguin-Jose Mantilla-Ramos, Ana-Sofía Hincapié-Casas, Annalisa Pascarella, Tarek Lajnef, Richard M. Leahy, Emily Coffey, Karim Jerbi, Véronique Boulenger

## Abstract

Low-frequency neural oscillations contribute to the parsing of continuous speech into linguistic units. Little is known however on the coupling of brain rhythms to higher-frequencies in speech such as fundamental frequency (F0) or pitch. Using magnetoencephalography, we investigated whole-brain cortical tracking of F0 while participants listened to sentences produced at normal rate or fast rate, where pitch naturally increases, and to artificially accelerated sentences, where F0 remains unchanged. Our results revealed significant brain-to-F0 coupling across all speech rates not only in right auditory but also in right parietal, insular, and pre- and postcentral regions, likely including the ventral larynx area. Importantly, the cortico-acoustic coupling peak frequency was higher for natural fast speech to reflect the corresponding F0 increase compared to normal rate and time-compressed speech. These findings demonstrate the engagement of an auditory-somato-motor network in F0 tracking, supporting its role in facilitating phonemic processing during the perception of naturally-produced speech.

## 1 Introduction

By tracking speech temporal structure, neural oscillations contribute to the parsing of the acoustic stream into hierarchically-organized linguistic constituents^1–4^. Substantial evidence from electro/magnetoencephalography (E/MEG) revealed phase-coupling of delta- (∼1-3 Hz) and theta-band (∼4-8 Hz) auditory cortical activity to the slow modulations of speech amplitude envelope, which mainly reflect the syllable rate^5–9^. Speech envelope has also been shown to modulate low (35-45 Hz) and high gamma (> 50 Hz) amplitude as well as theta phase/low gamma amplitude coupling^6,10–14^.

Besides syllabic structure, speech periodicity is another temporal cue of chief importance for speech perception and communication as it conveys segmental (voicing, manner of articulation) and suprasegmental (intonation, stress) information^15^. It results from the quasi-periodic vibration of the vocal folds during production and includes a representation of the speaker’s fundamental frequency (F0), which is the prime acoustic correlate of voice pitch. Only limited evidence exists regarding cortical tracking of F0 in naturally-produced speech. Many studies on pitch encoding measured the Frequency-Following Response (FFR), a sustained oscillatory response generally recorded by averaging responses to repeated syllables or tones that mimics stimulus periodicity and reflects precise auditory temporal processing^16–19^. Although the FFR recorded in EEG at the scalp is believed to originate primarily from the brainstem^20^, contributions from the thalamus and auditory cortex were also reported, with cortical activity peaking at the stimulus F0^21–28^. Brain coupling to F0 in continuous sound streams has been given less attention. The studies available so far focusing on neural encoding of the F0 in naturalistic speech showed responses in the auditory cortex at the F0 of isolated sentences or stories^29–32^. For instance, using a linear modelling approach in EEG, Van Canneyt and colleagues^33^ reported that the F0 of stories could be reconstructed from brain activity likely arising from the brainstem and the primary auditory cortex.

Here, we provide a novel contribution to this line of research by assessing whether neural oscillations, in the auditory cortex but also in other cortical regions, couple their oscillatory activity with the F0 of natural speech produced at different rates. We note that the term "coupling" is used here descriptively to refer to a statistical relationship between brain activity and the stimulus F0, without assuming whether this reflects the entrainment of endogenous oscillations or stimulus-driven phase-locked responses, a distinction that remains debated in the field^34^. Besides spectro-temporal modifications (e.g., shortening of acoustic cues, enhanced coarticulation)^35^, naturally speaking faster induces changes in the phonetic realization of F0. Acoustic studies in fact reported a shift in the frequency of the pitch range with increasing speech rate, namely decreased or increased pitch depending on speakers and languages^36,37^. We aimed to characterize how cortical F0 representation changes as a function of speech rate in natural speech.

Following earlier work showing that low-frequency cortical activity (e.g., 2-9 Hz) adjusts its coupling frequency to match the faster syllable rate^38,39^, we asked whether a similar frequency-shift occurs at higher frequencies to track the F0 changes accompanying speech rate increase. We note that the mechanisms underlying coupling at these different frequency scales likely differ: whereas theta/delta entrainment involves endogenous oscillatory dynamics, activity in the 77-97 Hz range examined here falls within the high-gamma band, which is generally thought to reflect local population firing rates rather than narrow-band oscillations^40,41^. Nonetheless, the question of whether brain-to-stimulus coupling at these frequencies exhibits a comparable rate-dependent frequency shift remains relevant and largely unexplored.

Using MEG, we examined brain coupling to the F0 of sentences naturally produced at a normal or fast rate, as well as of artificially accelerated (i.e. time-compressed) sentences. Given that time-compression reduces signal duration while keeping its spectral and pitch content intact^42^, time-compressed speech has the same F0 as normal rate speech despite a faster syllabic rate.

By contrast, F0 inherently increases in naturally-produced accelerated speech compared to both normal rate and time-compressed speech. Using Minimum Norm Estimation source reconstruction methods, we compared cortico-acoustic coupling and power modulations during sentence perception in the three speech rate conditions. In keeping with previous studies^7,38^, we analysed coupling in two frequency ranges centred on the mean F0 of our sentence material (77-84 Hz for normal rate/time-compressed sentences and 90-97 Hz for natural fast speech). In addition, a whole-brain analysis allowed us to investigate whether coupling to sentences’ F0 only occurs in the auditory cortex or whether it involves a more distributed cortical network, noting that previous work used region-of-interest based analyses, mostly in the primary auditory cortex^e.g.23,32^. We are aware of only one stereotactic EEG / MEG study showing FFR in a temporo-parieto-frontal cortical network in response to musical tones^43^.

Our results reveal significant cortico-acoustic coupling in frequency ranges that are specific to the sentences’ fundamental frequency. Coupling indeed occurred in the higher frequency band (90-97 Hz) for naturally-produced fast speech compared to normal rate speech, mirroring the natural increase in speech F0. No change in coupling frequency (77-84 Hz) was observed for time-compressed sentences which have similar F0 to normal rate sentences. Crucially, brain-to-F0 coupling was found in a right-lateralized network encompassing temporal, parietal, precentral and postcentral cortices, known to contribute to speech perception as well as to production. Such findings suggest that by adjusting to the rate of vocal fold vibration, neural oscillations in sensorimotor regions contribute to the encoding of pitch and of phonemic information conveyed by the periodicity of the speech signal.

## 2 Results

Twenty-four French native participants listened to natural normal rate (mean = 6.76 ± 0.57 syllables/s), natural fast rate (9.15 ± 0.60 syllables/s) and time-compressed sentences (9.15 ± 0.60 syllables/s) while their MEG brain activity was recorded. The mean F0 of the natural normal rate and time-compressed sentences was 81.3 Hz (SD 3.3) as calculated from the peaks of the power spectrum of the sentences using Python. It increased to 91.9 Hz (SD 9.8) for naturally accelerated sentences. Amplitude-modulated noise stimuli, generated with the envelopes of normal rate and fast rate stimuli, and which lack pitch and verbal information, were included as control conditions. We assessed neural tracking of changes in fundamental frequency by computing cortico-acoustic coupling between the acoustic signals and source-localized MEG time-series using phase-lag index (PLI) and, as a complementary approach, coherence measures (see Methods for more details about the selected coupling metrics). Because sentences’ F0 in the different speech rate conditions overlapped to some extent, we analysed speech-brain coupling in subsets of sentences for which the F0s of the normal and time-compressed conditions were most distant from the F0s of the fast condition (see Methods and Supplementary Figure S1 and Table S1). The F0 in these subsets peaked at 80 Hz for natural normal rate and time-compressed sentences, and at 93 Hz for natural fast rate sentences. We therefore conducted the coupling analysis in two frequency bands of interest centred on these values, respectively, 77-84 Hz and 90-97 Hz. For statistical analysis, PLI and coherence measures for actual stimulus encoding were compared with measures obtained on surrogate data using a shuffling procedure and correction for multiple comparisons using, respectively, cluster-based correction and FDR (see Methods).

### 2.1 Cortico-acoustic phase coupling shifts with speech F0

PLI analyses revealed significant phase-locking of neural oscillations to speech fundamental frequency, with a shift in coupling frequency that mirrored the changes in F0 following natural syllable rate acceleration (Figure 1). Cortico-acoustic coupling was found at 77-84 Hz for normal rate sentences mainly in bilateral Heschl’s, superior temporal, supramarginal and pre-/postcentral gyri as well as in the insula (see Supplementary Table S2 for more details). For the natural fast condition, PLI increased in the higher 90-97 Hz range in a right-lateralized network mostly including the same regions, namely Heschl’s, superior temporal, supramarginal and postcentral gyri (note from Figure 1A that coupling, albeit not significant, was also observed in the left hemisphere). As previously outlined, time-compressed sentences had the same F0 as normal rate sentences. This was well reflected in the brain by a significant PLI increase for this condition in the lower, normal rate pitch range only (77-84 Hz), in a similar network as the one found for normal rate speech. No significant phase-coupling was on the contrary observed for naturally accelerated sentences in this frequency range, stressing the specificity of the effect. Additional coherence analyses corroborated this pattern of results (see Figure 2). Indeed, coherence significantly increased bilaterally in the lower frequency range for normal rate and time-compressed sentences, but in the higher range for natural fast sentences. As expected, for the amplitude-modulated noise conditions, neither PLI nor coherence measures showed any significant cortico-acoustic coupling increase in any of the two frequency bands of interest (Figures 1 and 2).

**Figure 1.**
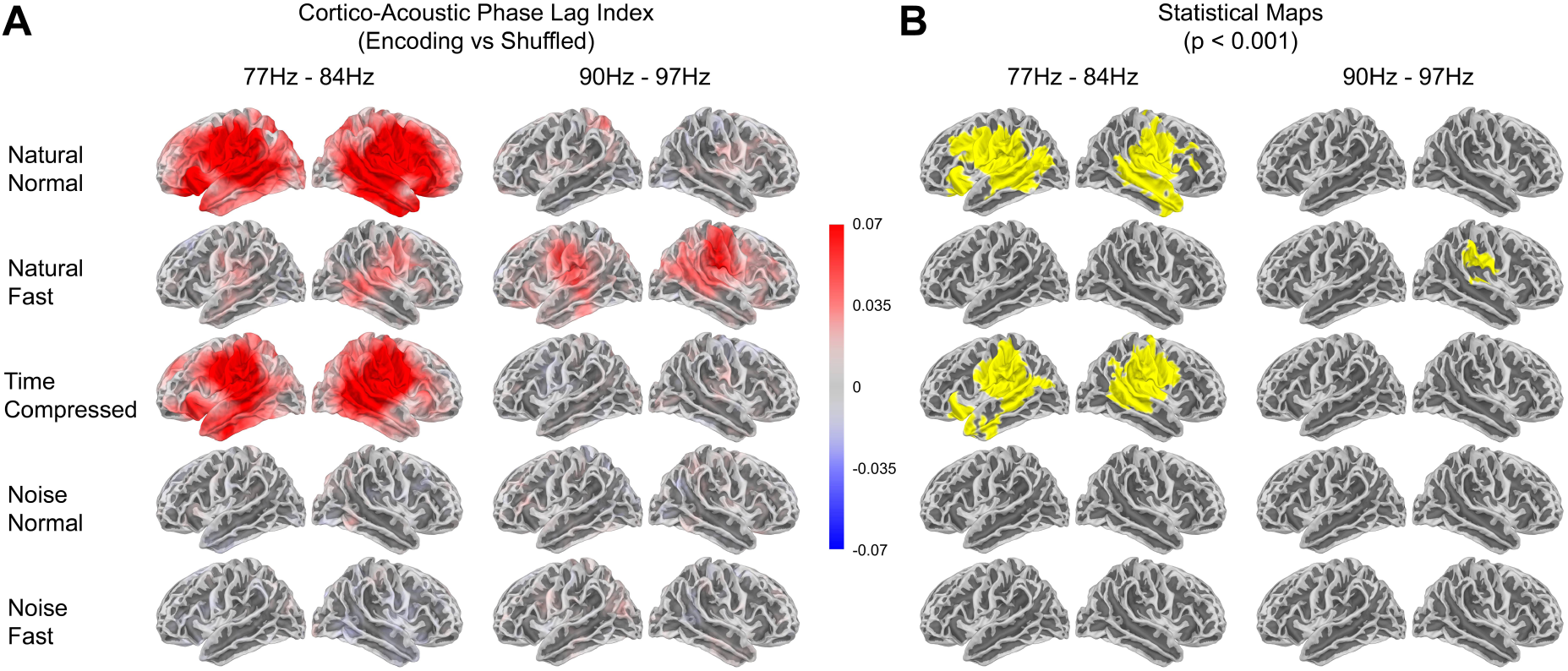
Cortical coupling to fundamental frequency in the frequency ranges encompassing the F0s of normal rate and time-compressed sentences (77-84 Hz) and of natural fast rate sentences (90-97 Hz). (A) Phase Lag Index (PLI) maps for coupling between the acoustic signal and brain oscillatory activity during stimulus encoding (active period), as compared to surrogate (i.e. shuffled) data. The values represent the difference in Fisher z-transformed PLI between the compared data. (B) Cluster-based statistical maps (corrected, α = .001, one-tailed) are shown for the different speech and noise conditions. Noise Normal and Noise Fast correspond to white noise stimuli modulated with the amplitude envelope of normal rate and fast rate sentences, respectively.

**Figure 2.**
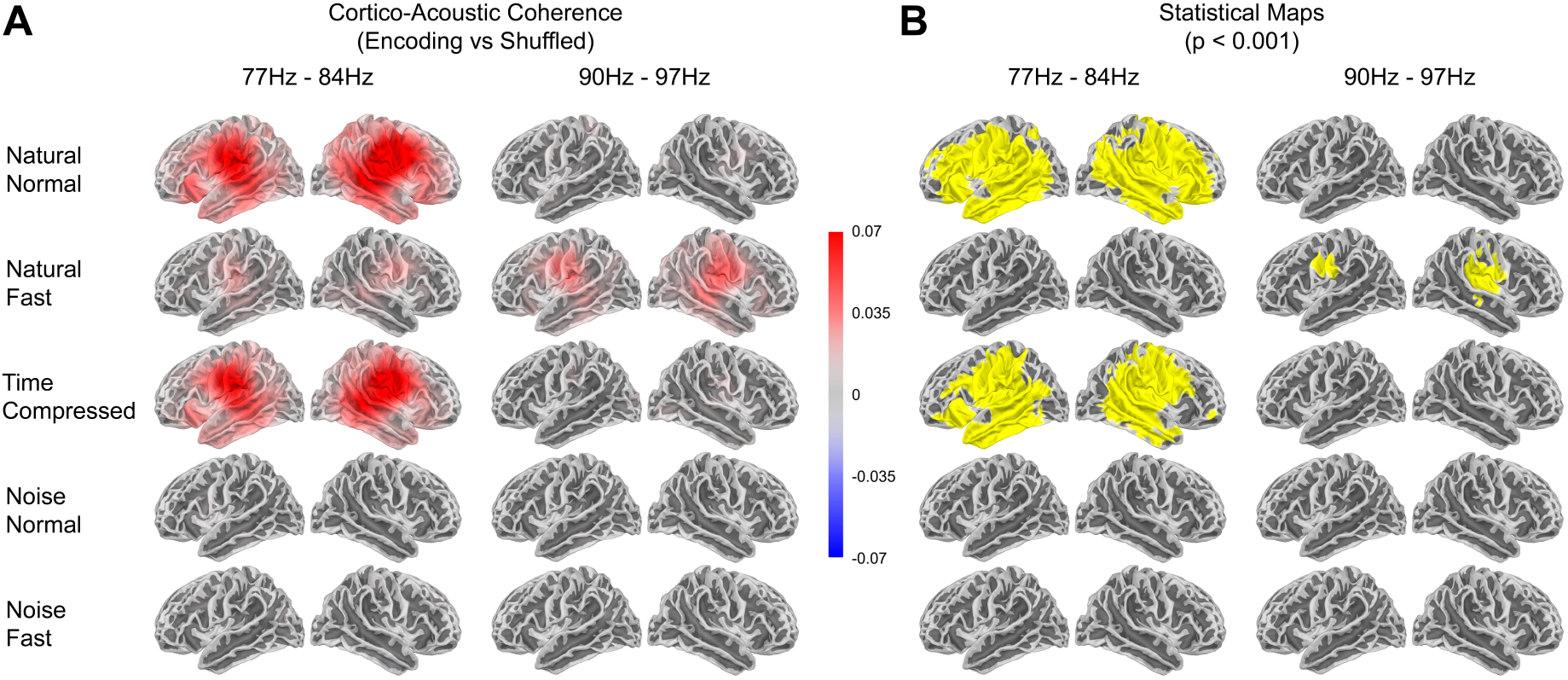
Cortico-acoustic coherence in the frequency ranges corresponding to the F0s in the normal rate and time-compressed conditions (77-84 Hz) and the natural fast rate condition (90-97 Hz). (A) Coherence maps for coupling between the acoustic signal and brain oscillatory activity during stimulus encoding (active period), as compared to surrogate (i.e. shuffled) data. The values represent the difference in Fisher z-transformed coherence between the compared data. (B) Statistical maps (FDR statistics, corrected, α = .001) are shown for the different speech and noise conditions. Noise stimuli modulated with the amplitude envelope of normal rate and fast rate sentences are referred to by Noise Normal and Noise Fast, respectively.

### 2.2 Coupling spectra identify top regions per parcellation

To more thoroughly assess the specific change in coupling frequency with F0 variation, we computed the coupling spectra for PLI and coherence across the three speech rate conditions, averaged over all participants and for each voxel. We then calculated the mean spectra separately over regions delimited by three different brain parcellations (see Methods). We identified the top regions in each parcellation with the highest spectral PLI peak across all speech conditions. For this analysis, we focused on the right hemisphere as it is where our PLI and coherence analyses above showed significant effects for all speech conditions. As displayed in Figure 3, this highlighted the Heschl’s gyrus (aparc parcellation), the first subzone of the postcentral gyrus (aparc_sub parcellation), and Brodmann area 43 corresponding to the subcentral area (PALS-B12 parcellation). Analyses confirmed that for both PLI and coherence, the coupling spectra in these regions peaked at the sentences’ F0, that is at ∼80 Hz for normal rate and time-compressed speech, with a shift at ∼90 Hz for naturally-accelerated speech (Figure 3 A, B, C for PLI and E, F, G for coherence). These peaks were not found when running the same analysis in the frontal pole which served as a control region (Figure 3 D, H). Note from Supplementary Figures S2, S3 and S4 that the three identified auditory, inferior postcentral and subcentral regions were found among the top 8 regions showing the highest PLI and coherence peaks at sentences’ F0, irrespective of the parcellation used. Note also that while power modulations might explain coherence peaks (Figure 3 E-M, F-N and G-O), this cannot be the case for the PLI curves as their calculation relies on phase measures rather than on the amplitude spectrum.

**Figure 3.**
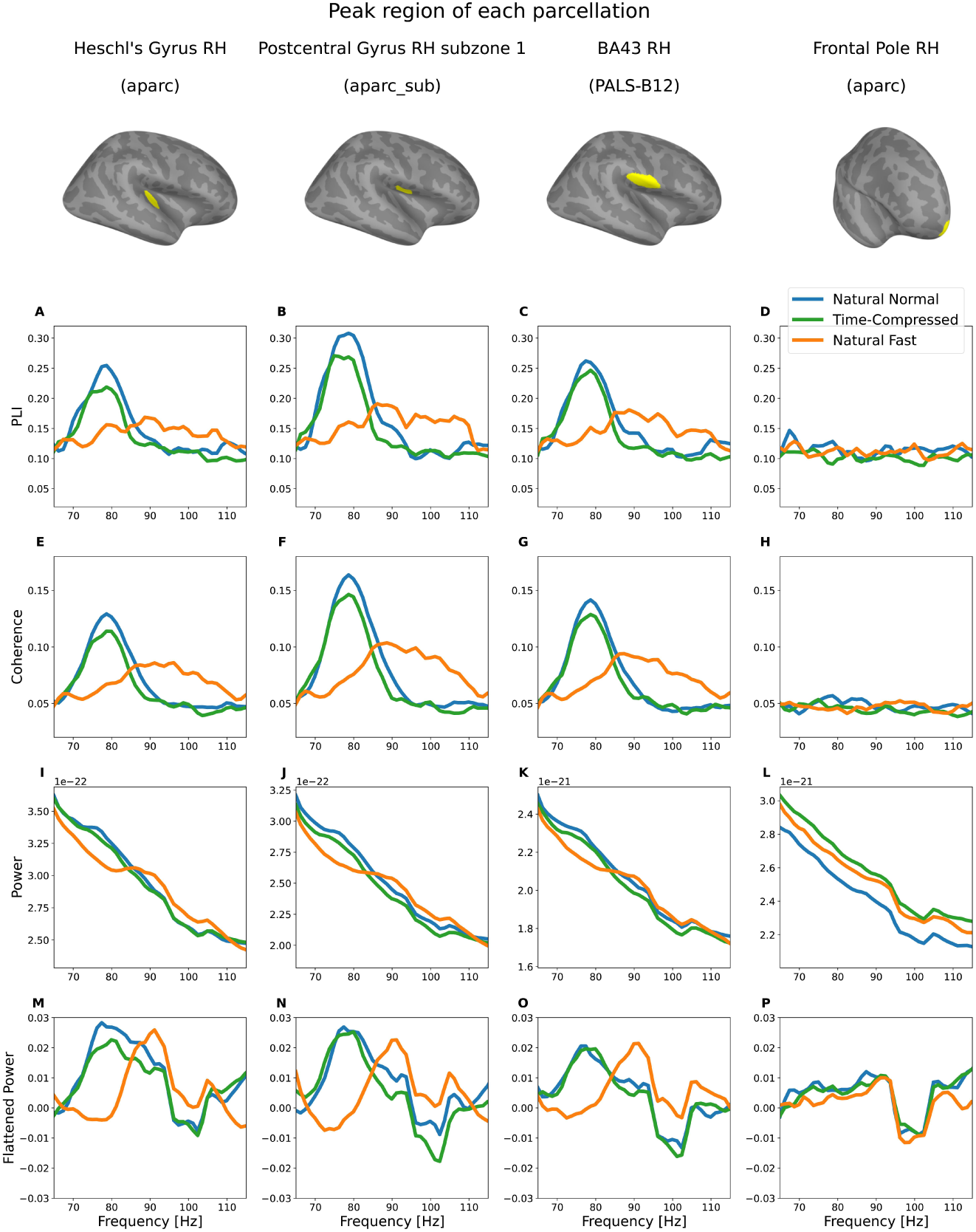
Coupling and power spectra for the three speech conditions in the top regions of the right hemisphere for each brain parcellation. The regions shown (one per column) correspond to the regions in each parcellation (respectively, aparc, aparc_sub and PALS-B12) with the highest peak of Phase Lag Index (PLI) coupling, which remains the same when evaluating coherence. The curves shown correspond to the average over subjects and voxels inside each region of PLI, coherence, power and flattened power, respectively. The coupling spectra were calculated between the acoustic signal and brain oscillatory activity during stimulus encoding (active period). For comparison, the same results are provided for the frontal pole as a control region. (A) to (D) show the PLI and (E) to (H) the coherence coupling spectra, respectively, for natural normal rate (in blue), time-compressed (in green), and natural fast rate speech (in orange). A detailed listing of the top 8 peak PLI regions for each of the three parcellations is provided in Supplementary Figures S2, S3 and S4. Notably, these top 8 peak regions include motor areas, further supporting F0 coupling beyond the auditory cortex. To evaluate the possible effects of power modulations on the coupling, (I) to (L) display the brain activity’s power spectrum during the encoding period and (M) to (P) show the same spectra but after correcting for the 1/f-like slope (using FOOOF^44^). The specific models used for the aperiodic fits of these curves can be seen in Supplementary Figure S5.

### 2.3 Power modulations mirror coupling findings

We finally examined whether brain coupling to speech F0 is also reflected in the amplitude of the cortical responses by analyzing source power modulations in each experimental condition and frequency band of interest. To this end, we contrasted power during the stimulus encoding period with power measured during the pre-stimulus baseline, using cluster-based statistical correction. Results showed a specific increase of power as a function of sentences’ F0 (Figure 4). In the normal and time-compressed conditions with F0 peaking around 80 Hz, brain activity significantly increased in the 77-84 Hz range in the right (and most of the time left for time-compressed speech) Heschl’s, superior temporal, supramarginal, insular and postcentral gyri. For natural fast rate sentences, the power increase was found at 90-97 Hz in a similar right-lateralized network also encompassing the precentral gyrus. No significant increase of power was found for amplitude-modulated noise stimuli in any of the frequency ranges.

**Figure 4.**
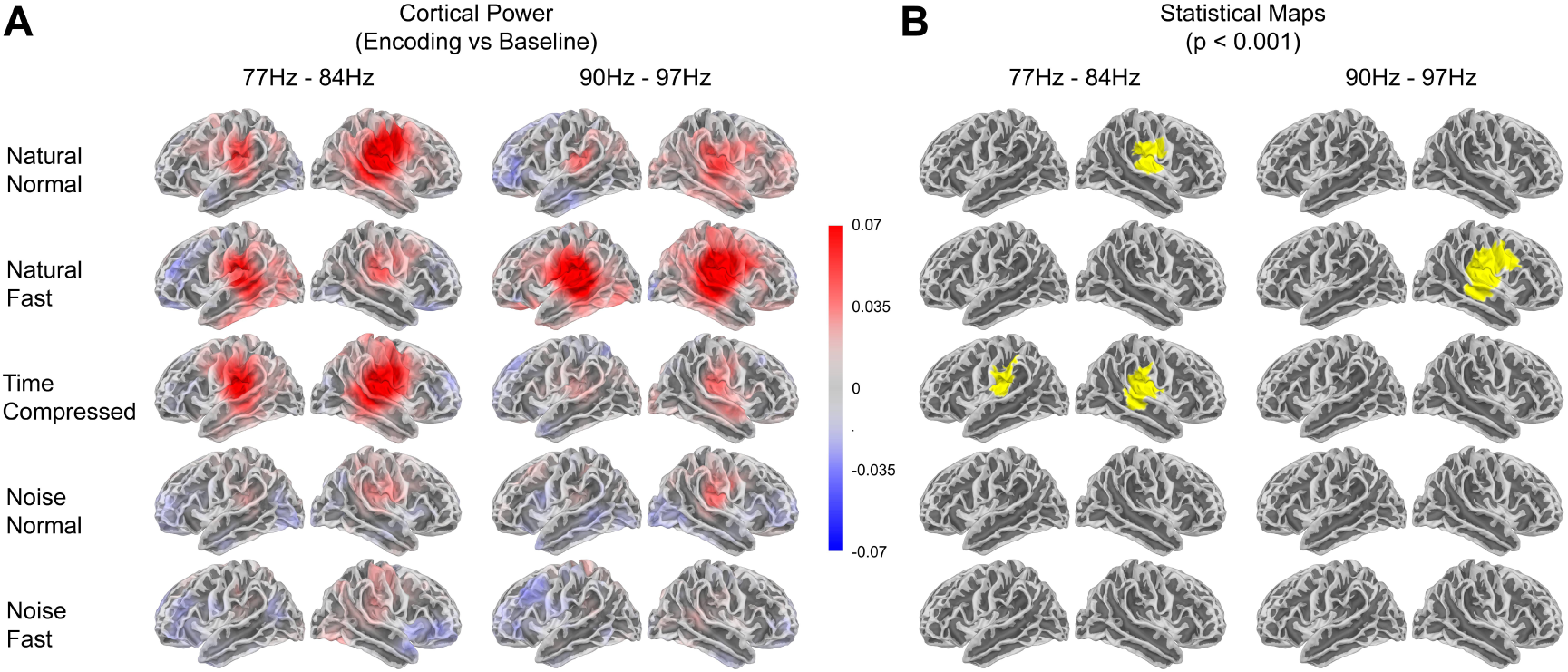
Power modulations in the frequency bands matching the F0s of natural normal rate and time-compressed sentences (77-84 Hz) and of natural fast rate sentences (90-97 Hz). (A) Power maps during the active period (i.e. stimulus encoding), as compared to the pre-stimulus baseline. The values represent the relative difference in Fisher z-transformed power between the compared data. (B) Cluster-based statistical maps (corrected, α = .001, two-tailed) are reported for the different speech and noise conditions.

## 3 Discussion

Our results revealed brain coupling to speech fundamental frequency, with a shift in coupling frequency and power increase that echoed the natural F0 increase with natural speech rate acceleration. Whereas phase-coupling (PLI), coherence and amplitude were enhanced at 77-84 Hz for normal rate and time-compressed speech (F0 peak at 80 Hz), coupling shifted to 90-97 Hz for naturally produced fast speech (F0 peak at 93 Hz). Crucially, this coupling was not restricted to the auditory cortex but encompassed a network distributed over sensorimotor regions. Overall, these findings therefore provide novel evidence that high-frequency cortical oscillations couple with the fundamental frequency of speech in the case of natural speech rate variations.

Coupling and power modulations to F0 were mainly found in the right or bilateral superior temporal (including the primary auditory cortex), supramarginal, pre-/postcentral gyri and insula, in line with previous work on pitch processing and voicing perception^45–49^. FFR studies also reported responses in the right or bilateral auditory cortex to syllables or continuous speech^23,26–28^. The lack of coupling to amplitude-modulated noise further suggests that phase-locking does not reflect responses to the temporal envelope of sentences but to their high-frequency carrier information. Importantly, our thorough whole-brain analysis using three different cortical parcellations highlighted the right Heschl’s gyrus, ventral postcentral gyrus and subcentral area (BA 43, at the inferior edge of the postcentral gyrus) as showing maximal coupling to sentences’ F0. Note the close spatial proximity of the latter two regions (14.7 mm based on a computation of distance between their centroids) which may correspond to the same somatosensory area. Each of these three regions furthermore showed up among the top eight regions with strong cortico-acoustic coupling across the three parcellations, indicating the reliability of the results. Other right-lateralized regions such as the insula and the supramarginal and precentral gyri also consistently coupled their oscillatory activity with speech F0, irrespective of the parcellation used. Interestingly, these regions are part of a dorsal network underlying speech auditory-motor integration^50,51^. The somatosensory cortex has also been advocated to contribute to speech perception by closely interacting with auditory regions^52^. In this regard, the subcentral gyrus (BA 43) identified in our study, owning both sensory and motor properties^53,54^, is seen as the ventral larynx motor area activated for the production of vocalizations and pitch control^55–59^. High-gamma activity in this region has been reported to increase before and during phoneme articulation^60^ and for voiced consonant production^61^, suggesting its role in the control of vocal fold movements. This area furthermore increases its connectivity with the auditory cortex during mere voicing perception^48^, and it can decode articulatory features of heard phonemes^62–64^. Our results concur with a functional role of this ventral somatosensory area in speech perception as it is one of the regions that best couple its high-frequency oscillatory activity with speech F0.

A dorsal larynx motor area, located in the precentral gyrus, is also thought to be involved in voicing and pitch production and perception^56,57,65–69^. In fact, and although this should be further tested, the precentral region coupling to speech F0 in our study may correspond to this dorsal motor region. This could reflect some simulation of the laryngeal gestures underlying pitch control during speech perception^70^. We also found reliable coupling to F0 in the supramarginal gyrus and insula. This agrees with the inferior parietal cortex acting as a sensorimotor interface to coordinate speech perception and production^51,71,72^ as well as with its role in phonological processing^73^. The insula is also assumed to contribute to auditory-motor speech integration^74^. It is indeed engaged in speech production and the control of laryngeal muscles and of F0 modulations^75–77^, but also in prosodic perception and auditory temporal processing^78–81^. In keeping with our work, high-gamma activity in the posterior insula was shown to track the F0 of a repetitive vowel similarly as in Heschl’s gyrus^82^. The insula moreover shares connections with auditory^83^ and motor regions, including the laryngeal motor cortex^84,85^. Altogether, our findings thus putatively suggest that besides the primary auditory cortex targeted by previous work^23,24,32,86^, a dorsal sensorimotor cortical network involved in speech auditory-motor integration syncs to the periodicity of vocal fold vibration in perceived speech.

Whether and how our findings relate to the cortical FFR to repetitive syllables^21,23,24,27,28^ is not straightforward. One difference is that the FFR is averaged over several hundreds of the same stimulus, whereas we assessed cortico-acoustic coupling and power modulations to a set of varying sentences. Still, the FFR is thought to indicate phase-locked neural activity to the stimulus F0. The different measures may therefore reflect at least some common processes. As previously mentioned, a FFR source in the right posterior auditory cortex has been described and would be involved in pitch computation^23,24^. Among the few studies that investigated F0 coupling to sentences or continuous speech, some focused on the brainstem^87–89^, while others examined high-gamma auditory cortex activity in light of the cortical FFR^26,30,32,33,90^, occasionally including regions extending slightly beyond primary auditory cortex (e.g., ref^26,30,32,33,90^); however, none systematically mapped F0 coupling across the whole brain at the source level. Crucially, by analyzing both coupling (using two measures, PLI and coherence) and power modulations at high frequencies across widespread cortical regions, we bring new support for brain coupling to F0 in natural speech that occurs well beyond the auditory cortex, in regions that are involved in auditory-somato-motor integration (in agreement with ^43^ for tones).

Although the increase in coherence we found might be explained, at least partly, by power increases at the same frequencies, this cannot hold for the phase-lag index which is insensitive to amplitude modulations. Cortical coupling to speech F0 may therefore capture the periodic nature of the phenomenon. Coffey and collaborators^86^ likewise proposed that their auditory cortex FFR entailed oscillatory entrainment since the brain response outlasted syllable offset (see also ^43^). Along with this view, oscillatory coupling to F0 in our study could reflect a predictive coding mechanism comparing speech input periodicity with internal temporal expectations. Such an interpretation, although still speculative, would support the idea that high-gamma activity mediates pitch-level prediction errors and their feedforward propagation up to higher integrative areas in order to update predictions^91–93^. Cortico-acoustic coupling to F0 in (pre)motor regions furthermore agrees with their known role in predictive coding, despite this being usually observed in lower (delta and beta) frequencies^94–96^. Further cross-frequency coupling and functional connectivity analyses between auditory and motor regions are needed to fully substantiate the predictive coding interpretation and further delineate the role of speech F0 coupling at the cortical level. As speech periodicity conveys segmental information, phase-locking to F0 might contribute to phoneme identification by encoding the articulatory feature of voicing or by extracting vowels’ formant structure. Another, not exclusive possibility is that it could serve more general communicative purposes, to track the relevant targets in an auditory scene^29^ and decrease sensitivity to environmental noise or competing speakers^86^.

Another question relates to the upper limit of F0 cortical tracking. In our study and previous work^23,27,86^, phase-locking was examined for typical male voices with F0 not exceeding 100-110 Hz. It has been proposed that the auditory cortex is not able, or hardly tracks frequencies beyond 100 Hz^20^, for pure tones^97,98^ or for syllables produced by males^21^ or females^33,90^. Some intracranial recordings still showed phase-locked responses up to 200 Hz^19,26,99,100^. In the present study, cortical tracking of natural fast speech was significant but less widespread than for normal rate and time-compressed speech, although the three conditions highlighted a comparable cortical network. This may relate to the general decrease in cortical frequency representation towards higher frequencies and/or the greater variance in F0 for rapidly-produced sentences. Part of the F0 distribution in this condition indeed lied beyond 110 Hz (up to ∼120 Hz) even after the selection procedure (Figure S1, panels B and D), which may have decreased the tracking capacity of cortical oscillations. Speech F0 therefore appears as an important parameter for the emergence of cortical tracking of speech periodicity, since phase-locking seems to decline with increasing F0. Alternatively, greater variability in natural fast speech F0 may have reduced statistical power, explaining the globally weaker effects in this condition. Further examining F0 tracking in different voices and under various speech rate or noisy conditions, will help better understanding this cortical contribution.

As in previous work in this domain, several critical aspects need to cautiously be taken into account when interpreting our speech-brain oscillatory coupling. An important interpretive caveat concerns whether the brain-to-F0 coupling we report reflects the entrainment of endogenous cortical oscillations to the stimulus periodicity, or rather stimulus-driven evoked responses phase-locked to each glottal pulse cycle^34^. Our data do not allow us to conclusively distinguish between these two accounts. However, several aspects of our results are more readily compatible with an oscillatory tracking mechanism. First, the PLI measure, which is insensitive to amplitude modulations, showed robust coupling, suggesting that the effect is not simply driven by repeated evoked power increases. Second, the frequency-specific shift in coupling that mirrors F0 changes across conditions, present for natural fast speech but absent for time-compressed speech which shares the same F0 as normal rate, points to a flexible mechanism that adjusts to stimulus periodicity rather than a fixed evoked response. Nonetheless, we acknowledge that these observations do not constitute definitive evidence for entrainment. This distinction is particularly relevant given that activity in the 77-97 Hz range falls within the high-gamma band, which is generally associated with local population firing rates rather than with narrow-band endogenous oscillations characteristic of theta/delta entrainment^40,41^. The frequency-specific coupling we observe may therefore reflect stimulus-driven synchronization of neural population activity to the periodicity of the glottal source, rather than the entrainment of intrinsic cortical rhythms. Future work using approaches such as post-stimulus analyses of sustained oscillatory activity^86^ will be needed to more firmly adjudicate between these accounts. Moreover, the reliability of the widespread network reported here would benefit from further investigation to rule out possible effects of spurious activity arising from the smoothing inherent to source reconstruction. Additionally, our coupling analyses were computed within narrow frequency bands centered on the mean F0 of each subset of sentences, which captures the brain’s sensitivity to the broad underlying periodicity of the stimuli but does not address whether cortical activity dynamically tracks the time-varying F0 contour within individual sentences (e.g., intonation patterns or local pitch fluctuations). Time-resolved approaches, such as temporal response functions or time-frequency coupling methods, would be needed to assess such fine-grained tracking and further characterize the cortical encoding of F0 dynamics.

Relatedly, our analysis was based on the mean F0 of each sentence, which captures the overall periodicity of the speech signal but does not account for within-sentence pitch variations related to prosodic contours (e.g., intonation, stress patterns). It is plausible that sentences with more stable F0 would yield stronger coupling than those with greater pitch variability, and that the higher F0 variance observed in the fast rate condition (SD 9.8 Hz vs. 3.3 Hz for normal rate) may have contributed to the weaker coupling effects in this condition. Future work could examine how within-sentence F0 variability modulates coupling strength, for instance by including F0 variance as a covariate in the analysis, and whether time-resolved methods can capture cortical tracking of prosodic pitch contours beyond the mean periodicity examined here. Importantly, while F0 is the primary acoustic correlate of pitch, prosody additionally encompasses variations in duration, intensity, and spectral quality, all of which are affected differently by natural acceleration and time compression. Natural fast speech alters the prosodic structure non-linearly, with changes in stress patterns and intonation that go beyond a simple shift in mean F0, whereas time-compressed speech preserves the original F0 contour but compresses its temporal realization. Our analyses focused on coupling to the mean F0 periodicity and were not designed to capture these broader prosodic dimensions. As noted above, the greater F0 variability in the fast rate condition (SD 9.8 Hz vs. 3.3 Hz for normal rate) may have contributed to the weaker coupling observed for this condition. A promising future direction would be to model F0 continuously on a trial-by-trial basis, for instance using regression approaches that test whether neural coupling scales with F0 rather than relying on categorical frequency bands, and to examine how within-sentence prosodic variations modulate cortical tracking.

In conclusion, our study suggests that high-frequency cortical activity couples with the fundamental frequency (F0) of speech, with a shift in coupling frequency that mirrors the F0 variation accompanying natural speech rate acceleration. Using magnetoencephalography (MEG), we identified significant brain-to-speech F0 coupling across a network extending beyond the auditory cortex to include the right auditory, inferior parietal, insular, and pre- and postcentral regions, involving a dynamic auditory-somato-motor network in the encoding of speech. These findings advance our understanding of how high-frequency cortical activity contributes to the encoding of speech fundamental frequency, and point to a complex interplay between auditory and motor systems in speech perception. Future work using time-resolved approaches and diverse speaker profiles will help further characterize these cortical coupling mechanisms and their functional significance for language processing.

## 4 Methods

### 4.1 Participants and Ethics

Twenty-four (14 females) participants aged between 18 and 45 years old (mean = 23 years old) volunteered for the study. All were monolingual French native speakers and right-handed (mean score at the Edinburgh handedness inventory^101^ = 94). None of them reported hearing or language problems nor history of neurological or psychiatric disorders. The protocol conformed to the Declaration of Helsinki and was approved by the local ethical committee (Comité de Protection des Personnes Lyon Sud-Est II; ID RCB: 2012-A00857-36). Written informed consent was obtained for all participants before the experiment, and they were compensated for their participation.

### 4.2 Stimuli

A set of 288 meaningful sentences (7-9 words) with the same syntactic structure (e.g., “Le guide organise une excursion dans le désert” / *The guide organizes a trip to the desert*) served as stimuli. A French native male professional theatre actor recorded the stimuli in a sound-attenuated booth using ROC*me*! software ^102^. Each sentence was first displayed on screen for the actor to silently read before producing it aloud at his normal speech rate. He then produced the same sentence as fast as possible while remaining intelligible, with no externally imposed target rate. Recordings were sampled at 44.1 kHz (mono, 16 bits). The mean syllable rate of the sentences (i.e. number of produced syllables divided by sentence duration) as calculated with Praat^103^, was 6.76 syllables/s (SD 0.57) for the natural normal rate condition, while it increased to 9.15 syllables/s (SD 0.60) in the natural fast condition. Time compression, which is a re-synthesis process only affecting the temporal structure of the stimuli, was then applied to normal rate sentences by digitally shortening them with a PSOLA algorithm (Pitch Synchronous Overlap and Add^104^). Compression rates were computed for each sentence, namely each time-compressed sentence was matched in terms of syllable rate to its natural fast equivalent. Therefore, time-compressed sentences had a mean syllable rate of 9.15 syllables/s (SD 0.60) as well. Mean sentence duration was 1.67 s (SD 0.16) for natural normal rate, 1.23s (SD 0.14) for natural fast rate, and 1.24 s (SD 0.14) for time-compressed sentences, corresponding to a total stimulus duration of approximately 610 s, 449 s, and 449 s per condition, respectively. For the total of 864 sound files (288×3 speech rate variants), we applied an 80 Hz high-pass filter and smoothed sentences’ amplitude envelope initially and finally. The intensity of the sound files was finally peak normalized.

Computation of the power spectrum in the 64-128 Hz range across sentences in the three speech rate conditions using Python showed peak frequencies at 81.6 Hz (SD 3.2) and 81 Hz (SD 3.3) for the normal rate and time-compressed conditions respectively, and at 91.9 Hz (SD 9.8) for the fast rate condition (see Figure S5 in the supplementary material). These peak frequencies are consistent with the computation of the median pitch values with Praat.

To assess whether brain-to-F0 coupling occurs with the high-frequency carrier information in speech and not with the modulations of the amplitude envelope, we generated amplitude-modulated noise stimuli as control conditions. A total of 576 control stimuli were created by modulating Gaussian white noise (lacking linguistic content) with the amplitude envelope of each of the 288 natural normal rate and 288 natural fast rate sentences. We also added forty-eight filler sentences (different from the experimental stimuli but following the same syntactic structure) in which beep-sounds were inserted at different possible time-points (i.e. 200, 600 or 800 ms from onset) in the sentences. These filler sentences were used to encourage participants to pay attention to the auditory stimuli during the experiment (see Experimental procedure). The stimuli were distributed across two experimental lists, each including 288 sentences (96 for each speech rate condition), 192 amplitude-modulated noise stimuli (96 modulated with the normal rate envelope and 96 with the fast rate envelope) and 48 filler sentences, for a total of 528 stimuli per list. Each participant was presented with one list, in which each sentence appeared in only one of the three rate conditions (i.e., no sentence was repeated within a participant). The assignment of sentences to rate conditions was rotated across the two lists, such that sentences appearing in both lists were assigned to different rate conditions in each.

### 4.3 Experimental procedure

While comfortably sitting in a sound-attenuated, magnetically-shielded recording room with a screen in front of them, participants were presented binaurally with stimuli through air-conducting tubes with foam insert earphones (Etymotic ER2 and ER3). Prior to the MEG recording, their auditory detection thresholds were determined for each ear with a pure tone. The level was then adjusted in order to present stimuli at 50 dB Sensation Level (SL) with a central position (stereo) with respect to the participant’s head.

Each trial of the experiment ran as follows: after a fixation cross displayed at the centre of the screen for 1500 ms, an auditory stimulus (sentence or amplitude-modulated noise) was presented, with the fixation cross remaining on the screen. Participants were instructed to attentively listen to all stimuli while looking at the fixation cross and to press a button (response button Neuroscan, Pantev) with their left index finger as quickly as possible whenever they detected beep-sounds in (filler) sentences. The next trial started after an inter-trial interval (grey screen) of 1250 ms. Participants detected 100% of the filler trials, which were excluded from subsequent analysis. Each participant listened to the stimuli from one of the two experimental lists. The total of 528 stimuli were pseudo-randomly divided into 8 blocks (66 stimuli each) separated by short breaks to allow the participants to rest. Participants were familiarized with the stimuli and task in a training phase (5 sentences different from the experimental stimuli). To further maintain their attention throughout the experiment, they were informed that questions on the heard sentences would be asked at the end of the experiment. More precisely, they had to indicate, by ticking a yes/no box on a paper sheet, whether eight proposed sentences (three of which were not experimental stimuli) were presented during the MEG experiment. The experiment was run with Presentation software (Neurobehavioral Systems).

### 4.4 MEG data acquisition and analysis

We recorded neuromagnetic brain activity of the 24 participants using a 275-channel whole-head MEG system (CTF OMEGA 275, Canada) at 1200 Hz sampling rate. Head position within the MEG helmet was monitored with three fiducial coils (nasion, left and right pre-auricular points) placed on the participant’s head. Horizontal and vertical eye movements were also recorded with four electrooculography (EOG) electrodes. We recorded reference head position before each of the 8 experimental blocks and tracked head movements throughout the experiment using continuous head position identification (HPI).

Preprocessing of the MEG data was done with custom written Matlab scripts (Mathworks Inc., MA, USA) and the Fieldtrip toolbox^105^. Subsequent coupling analyses were carried out in Python using custom scripts and the MNE-Python library^106^. In the following we describe the processing of 1) speech recordings, 2) MEG data, 3) Magnetic Resonance Imaging (MRI) data, and the analyses of 4) source localization, power and coupling, and 5) coupling maps and statistics.

#### 4.4.1 Audio Recordings (reference signals)

To examine the neural tracking of speech F0, we used the speech signal as the reference. Inspection of the F0 distributions and power spectral densities revealed some spectral (including F0s) overlap between sentences in the different speech rate conditions (see Supplementary Figure S1 A and C). To test our hypothesis that brain oscillatory activity adjusts to F0 increase with speech rate acceleration, we optimized the distinction between conditions in terms of F0s by selecting a proportion of sentences in the natural normal rate and time-compressed conditions with the lowest mean F0s (71% and 77% of the sentences on average per participant, respectively). Similarly, we selected sentences in the natural fast rate condition with the highest mean F0s (∼62% on average per participant). To do this, we collected sentences from the normal rate/time-compressed and fast rate conditions starting from opposite sides of the spectrum, thus maximizing the spectral distance between these two subsets while keeping an acceptable number of sentences in each condition (see Supplementary Table S1). This resulted in normal rate and time-compressed sentences with the lowest F0s (i.e. below 83 Hz) and a F0 peak at 80 Hz, and natural fast rate sentences with the highest F0s (i.e. above 87 Hz) peaking at 93 Hz (see Supplementary Figure S1 B and D). We analyzed brain-to-speech coupling in two frequency ranges centered on the F0 peak values (across sentences) of the two subsets of sentences (with 7Hz bandwidth), namely 77-84 Hz and 90-97 Hz. We note that this sentence selection procedure transforms a continuous acoustic variable into a categorical one, and that excluding sentences with overlapping F0 reduces statistical power. However, this step was necessary to ensure sufficient spectral separation between conditions for a meaningful test of frequency-specific coupling. Importantly, the selection retained a majority of sentences in each condition, and the coupling spectra computed over the full frequency range (Figure 3) confirm that the observed peaks are not artifacts of this selection. For this coupling analysis, we selected speech segments between 200 ms and 1000 ms post-stimulus onset, as the shortest sentences in the natural fast and time-compressed conditions were ∼1 s long (this time-window for speech segments equals the time-window of interest of the functional MEG data, see next section).

#### 4.4.2 MEG data

The MEG continuous data were segmented into 3 s periods (from 1 s before to 2 s after stimulus onset). A standard semi-automatic procedure available in the Fieldtrip toolbox was used to reject data segments contaminated by eye blinks, heartbeat and muscle artefacts. To remove the power line artefact and its harmonics notch-filtering was applied at 50, 100 and 150 Hz. We then detected and rejected EOG, jumps and muscle artifacts and visually double-checked the trials. Finally, an Independent Component Analysis (ICA) was used to correct for ECG artifacts. For each trial, the time interval of interest was set between 200 ms and 1000 ms after stimulus onset, while the baseline extended from 1000 ms to 200 ms before onset. The cleaned and segmented data was then imported to Python to perform the coupling analysis.

#### 4.4.3 MRI data

T1-weighted structural MRIs (MRI 1.5 T, Siemens AvantoFit) were acquired for 23 of the 24 participants after the MEG study. Alignment of each MRI to the MEG coordinate system was performed with the localization coils marked on the MRIs and the interactive alignment interface available in MNE. Each MRI was segmented to compute each participant’s head model using the single-shell as volume conductor model^107^. To carry out group analyses in a common space, we first spatially normalized a template anatomical grid from the Montreal Neurological Institute (MNI) common coordinate system. MRI grids of each participant were then aligned to this normalized template (this common grid has 5124 vertices). We finally computed the cortical source space modelling for each participant using the individual MRI volume, but with the surface grid based on the MNI template. This allowed transforming the results to the same brain template for all participants for easier comparison.

#### 4.4.4 Source localization, power and cortico-acoustic coupling estimation

We estimated source power and cortico-acoustic coupling given source reconstructions calculated using Minimum Norm Estimation^108–110^. Before computing the coupling, we resampled the speech signal to 1200 Hz to match the sampling frequency of the MEG data. In this way, we could suitably assess the neural tracking of F0 by computing the coupling between auditory signals and cortical activity using Phase Lag Index and coherence measures. The Phase Lag Index provides information regarding the coupling of two signals with respect to the sign of their phase difference^112^. It has the advantage of being insensitive to the amplitude of the signals and moreover, neglecting zero-phase coupling that commonly arise because of volume conduction. It can be mathematically defined as:

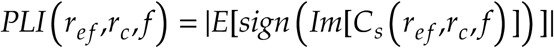

where ***r_ef_*** is the reference signal, namely the acoustic signal here; ***r_c_*** is the brain signal at each vertex of the anatomical grid; ***f*** is the frequency bin and ***Cs*** is the Cross Spectral Density (CSD) matrix. The operator E[x] refers to the average over trials, |x| is the absolute value, sign(x) returns the sign of the value, and Im[x] gives the imaginary part of a complex number.

Coherence refers to the linear correlation between two signals as a function of frequency. It can be mathematically defined as:

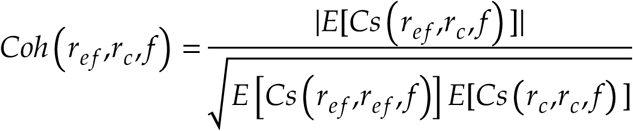

where the notation is the same as the one used in the previous equation.

We computed PLI and coherence using the MNE-Connectivity package within the MNE-Python ecosystem^106^. Specifically, the CSD matrix was computed using the multitaper Fast Fourier Transform (FFT) from 64 to 128 Hz, with 7 DPSS tapers (bandwidth ≈ 10 Hz) applied to 0.8 s segments (961 time points), yielding a frequency resolution of approximately 1.25 Hz. Cortico-acoustic coupling was calculated for the active period (200 to 1000 ms), the baseline (−1000 to −200 ms), and for shuffled data, the last two serving as control conditions for statistical analyses for power and coupling, respectively. In the case of surrogate data, we randomly shuffled the samples of the audio signals before computing cortico-acoustic coupling (i.e. the structure of the audio signal was lost and consequently did not match its MEG counterpart for the coupling computation). We repeated this procedure 100 times and averaged coupling across iterations for each participant, and each experimental condition and frequency bin. For both active, baseline and shuffled data, the couplings (and power) were averaged across the frequency bins inside each band of interest.

#### 4.4.5 Cortico-acoustic coupling maps, power maps and statistical analyses

The average coupling and power maps used for visualization in the Results section were obtained by first applying the Fisher transform of the values. Then we computed the difference between the active period and a set of control data (shuffled audio for coupling, and baseline for power) for each participant and condition. In the case of power, the relative difference was used. The values were then averaged across participants for each experimental condition and subsequently mapped to the brain plots. Group statistical analysis was conducted on the raw coupling and power values (without the Fisher transform) for the 23 participants with individual MRIs. For PLI and power, we applied cluster-based permutation tests with 1024 permutations^113^. For coherence, we performed an FDR corrected t-test. The tests are one-tailed for cortico-acoustic coupling analysis and two-tailed for power analysis. All tests use a statistical significance threshold of 0.001. We used the Desikan-Killiany atlas to identify the regions detected as significant in the statistical analysis^114^. The top 10 regions with the highest mean t-value based on the PLI statistical analysis for each frequency range, condition, and hemisphere can be found in Supplementary Table S2.

To inspect more closely the changes in coupling frequency with F0 variation, we additionally computed - for each of the three speech rate conditions - the coupling spectra during the encoding period, both for PLI and coherence, across a wide range of frequencies (64-128 Hz) and averaged through participants. This was done on all voxels and then averaged into the regions of the Desikan-Killiany atlas^114^ (aparc parcellation). We simultaneously explored two other parcellations: aparc_sub^115^ (corresponding to subdivisions of the regions of the aparc parcellation) and PALS-B12^116^ (which delineates Brodmann areas). Then, the top 8 regions in the right hemisphere with the highest PLI peak independent of the speech rate conditions were identified for each of the three parcellations. These regions are detailed in Figures S2, S3 and S4. We further compared the top region of each parcellation with the frontal pole, selected as the control region since it was not expected to show any F0 response (see Figure 3; see also^23^). Additionally, to assess the potential effects of power modulations, we computed the power spectra of the encoding brain signals at these locations in a similar manner. To better compare the coupling spectra to the oscillatory peaks in the brain power, we removed the aperiodic component using the FOOOF (specparam) methodology^44^. The details of the model that fits each of the spectra involved in Figure 3 can be seen in Supplementary Figure S5.

## Data and Code Availability

Custom analysis code used in this study is openly available at https://doi.org/10.5281/zenodo.19637465. The data that support the findings of this study are available from the corresponding authors upon reasonable request. Access requires a formal data sharing agreement, approval from the requesting researcher’s local ethics committee, and submission of a formal project outline. Requesters are expected to cite the original data source and the present manuscript.

## CRediT authorship contribution statement

Yorguin-Jose Mantilla-Ramos: Data curation, Formal analysis, Methodology, Software, Visualization, Writing – original draft. Ana-Sofía Hincapié-Casas: Data curation, Formal analysis, Methodology, Software. Annalisa Pascarella: Methodology. Tarek Lajnef: Methodology. Richard M. Leahy: Methodology. Emily Coffey: Writing – review & editing. Karim Jerbi: Conceptualization, Methodology, Software, Validation, Resources, Writing – original draft, Supervision, Project administration, Funding acquisition. Véronique Boulenger: Conceptualization, Methodology, Validation, Investigation, Resources, Writing – original draft, Supervision, Project administration, Funding acquisition.

## Acknowledgements

We thank Damien Gouy for recording the stimuli, Emmanuel Ferragne for his help with Praat software, and Mélanie Canault for discussions on articulatory phonetics. AI-assisted language tools were used during the preparation of this manuscript to improve text readability. All authors reviewed and edited the output and take full responsibility for the content of the publication.

