## Supplementary for "Hitting the right pitch: Cortical tracking of speech fundamental frequency in auditory and somatomotor regions"

### Supplementary Material

Content:

Table S1-S2

Figures S1-S5

| Condition | Lowest % (and number) of sentences kept across participants | Mean % (and number) of sentences kept across participants | Highest % (and number) of sentences kept across participants |
|---|---|---|---|
| Normal rate | 66.67% (37) | 70.95% (53.04) | 79.63% (64) |
| Time-Compressed | 71.25% (42) | 77.41% (58.39) | 83.13% (69) |
| Fast rate | 55.88% (35) | 62.10% (46.65) | 70.27% (57) |

**Table S1: Mean percentage and percentage range of included sentences across participants by speech rate condition, after selecting sentences to reduce F0 overlap between subsets of sentences in the three conditions.** For each MEG recording (corresponding to a participant and a condition), sentences were selected to reduce the F0 overlap between the Fast rate condition and the other two conditions (Normal rate and Time-Compressed). After this procedure, the number and percentage of included sentences within that MEG recording were calculated. Subsequently, the inclusion values were characterized by mean and range (lowest, highest) across participants within a condition. None of the participants were excluded after this selection procedure. The minimum % of included sentences is ~56% for one participant in the fast rate condition. Nevertheless, for this condition, the overall mean inclusion percentage is ~62% (with a maximum of ~70%). All conditions have around 3% standard deviation of the inclusion percentage. The values in brackets represent the number of sentences corresponding to the given percentage. For the range, these are integer numbers as they represent a specific participant. For the mean, fractional values appear as these represent the average value of included sentences across participants.

| Condition | 77-84 Hz | | | | | | | | 90-97 Hz | | | | | | | |
|---|---|---|---|---|---|---|---|---|---|---|---|---|---|---|---|---|
| | Left Hemisphere | | | | Right Hemisphere | | | | Left Hemisphere | | | | Right Hemisphere | | | |
| | Region | Mean t-value | SD | Voxel Count | Region | Mean t-value | SD | Voxel Count | Region | Mean t-value | SD | Voxel Count | Region | Mean t-value | SD | Voxel Count |
| Normal | Transverse Temporal Gyrus | 5.95 | 0.42 | 17 | Posterior Cingulate Cortex | 5.05 | 0.98 | 41 | None | | | | None | | | |
| | Superior Temporal Gyrus | 4.95 | 0.83 | 79 | Transverse Temporal Gyrus | 4.79 | 0.35 | 10 | | | | | | | | |
| | Banks of the Superior Temporal Sulcus | 4.85 | 0.58 | 35 | Insular Cortex | 4.56 | 0.67 | 52 | | | | | | | | |
| | Postcentral Gyrus | 4.75 | 0.66 | 72 | Paracentral Lobule | 4.53 | 0.62 | 30 | | | | | | | | |
| | Insular Cortex | 4.72 | 0.88 | 62 | Supramarginal Gyrus | 4.49 | 0.63 | 45 | | | | | | | | |
| | Inferior Parietal Lobule | 4.71 | 0.83 | 33 | Postcentral Gyrus | 4.33 | 0.45 | 73 | | | | | | | | |
| | Caudal Middle Frontal Gyrus | 4.40 | 0.45 | 18 | Superior Temporal Gyrus | 4.08 | 0.39 | 93 | | | | | | | | |
| | Supramarginal Gyrus | 4.29 | 0.61 | 82 | Precentral Gyrus | 4.02 | 0.41 | 114 | | | | | | | | |
| | Precentral Gyrus | 4.28 | 0.51 | 75 | Banks of the Superior Temporal Sulcus | 3.86 | 0.22 | 30 | | | | | | | | |
| | Middle Temporal Gyrus | 4.18 | 0.45 | 36 | Inferior Parietal Lobule | 3.79 | 0.17 | 5 | | | | | | | | |
| | Left Hemisphere | | | | Right Hemisphere | | | | Left Hemisphere | | | | Right Hemisphere | | | |
| | Region | Mean t-value | SD | Voxel Count | Region | Mean t-value | SD | Voxel Count | Region | Mean t-value | SD | Voxel Count | Region | Mean t-value | SD | Voxel Count |
| Fast | None | | | | None | | | | None | | | | Supramarginal Cortex | 4.08 | 0.40 | 64 |
| | | | | | | | | | | | | | Transverse Temporal Cortex | 3.91 | 0.28 | 6 |
| | | | | | | | | | | | | | Postcentral Cortex | 3.91 | 0.30 | 21 |
| | | | | | | | | | | | | | Superior Temporal Cortex | 3.89 | 0.24 | 13 |
| | Left Hemisphere | | | | Right Hemisphere | | | | Left Hemisphere | | | | Right Hemisphere | | | |
| | Region | Mean t-value | SD | Voxel Count | Region | Mean t-value | SD | Voxel Count | Region | Mean t-value | SD | Voxel Count | Region | Mean t-value | SD | Voxel Count |
| Time Compressed | Transverse Temporal Gyrus | 4.91 | 0.51 | 17 | Transverse Temporal Gyrus | 5.11 | 0.43 | 10 | None | | | | None | | | |
| | Supramarginal Gyrus | 4.77 | 0.60 | 98 | Posterior Cingulate Cortex | 4.95 | 0.86 | 38 | | | | | | | | |
| | Postcentral Gyrus | 4.70 | 0.65 | 101 | Supramarginal Gyrus | 4.95 | 0.87 | 104 | | | | | | | | |
| | Insular Cortex | 4.62 | 0.66 | 42 | Postcentral Gyrus | 4.71 | 0.66 | 89 | | | | | | | | |
| | Posterior Cingulate Cortex | 4.60 | 0.52 | 34 | Insular Cortex | 4.61 | 0.72 | 45 | | | | | | | | |
| | Superior Temporal Gyrus | 4.34 | 0.52 | 72 | Superior Temporal Gyrus | 4.51 | 0.50 | 59 | | | | | | | | |
| | Precentral Gyrus | 4.15 | 0.46 | 42 | Paracentral Lobule | 4.25 | 0.50 | 33 | | | | | | | | |
| | Temporal Pole | 4.14 | 0.34 | 9 | Banks of the Superior Temporal Sulcus | 4.22 | 0.30 | 36 | | | | | | | | |
| | Banks of the Superior Temporal Sulcus | 4.12 | 0.26 | 29 | Caudal Anterior Cingulate Cortex | 4.14 | 0.21 | 9 | | | | | | | | |
| | Lateral Orbitofrontal Cortex | 4.11 | 0.40 | 40 | Precentral Gyrus | 4.12 | 0.44 | 106 | | | | | | | | |

**Table S2: Top 10 statistically significant regions from the cluster-based analysis of the PLI for each condition, frequency range and hemisphere.** This represents a more tabular view of Figure 1. For each condition (Normal, Fast, Time-Compressed), frequency range (77-84 Hz and 90-97Hz) and hemisphere (left and right), the statistically significant regions with the highest mean t-value (averaged over the voxels of the region) are shown. Each region is also described by its number of voxels (voxel count) and the standard deviation of the t-values (SD).

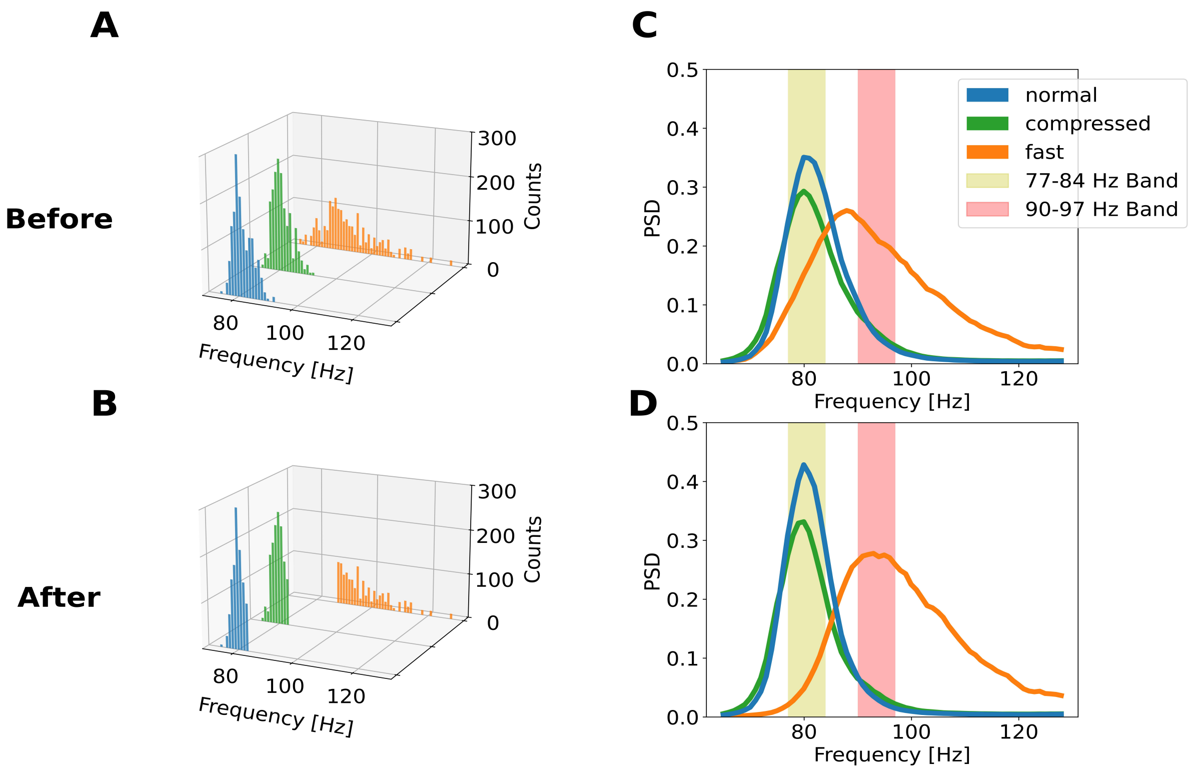


**Figure S1: Optimizing sentence selection (normal rate vs. fast rate) in order to minimize F0 overlap across speech conditions.** Sentences with the lowest F0s (<83 Hz) in the normal rate/time-compressed conditions and sentences with the highest F0s (>87 Hz) in the fast rate condition were selected. In this way, the spectral distance between the two subsets of stimuli was maximized. Sentences were chosen in such a way that at least 60% of them were retained in each condition, irrespective of which participant they correspond to. This resulted in a selection of, on average, 71% of the normal rate sentences, 77% of the time-compressed sentences, and 62% of the fast rate sentences (see Table S1 in the supplementary material for the percentage of sentence selection across participants). (A) shows the original distribution of F0s which exhibits the overlap of F0 across the sets of stimuli (i.e. before the selection procedure). (B) displays the distribution of F0s after the selection procedure was applied. Note the absence of F0-overlap between the two subsets. (C) Mean spectral density of the audio spectrum per condition prior to reducing the overlap (corresponding to panel A). The highest portion of the curves for normal rate and time-compressed sentences overlaps with the lowest portion and almost the peak of the curve for fast rate sentences. (D) Same as (C) but showing the mean power spectral density after the stimulus selection procedure. Notice the right shift of the peak of audio spectra for fast rate sentences and the reduced overlap of its lowest portion with the highest portion of normal rate and time-compressed curves. The two bands of interest that we used for the coupling analysis (77-84 Hz and 90-97 Hz) are shaded, respectively, in gold and red in panels (C) and (D).

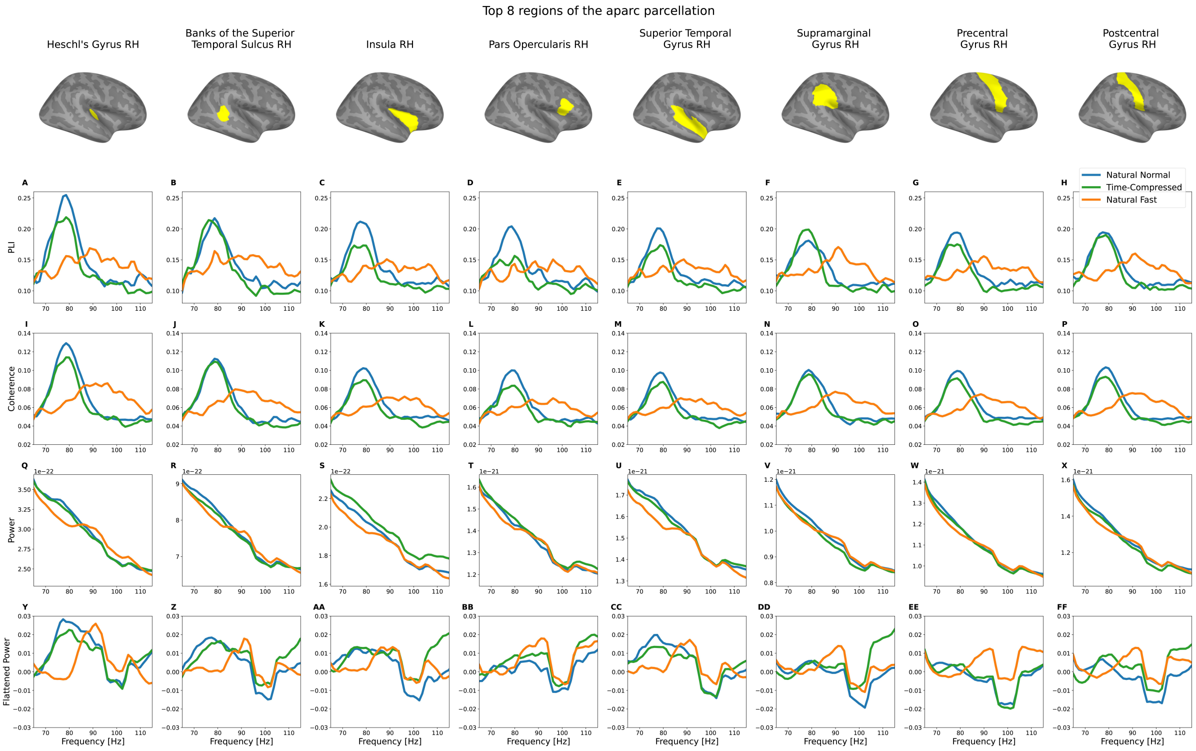


**Figure S2: Top 8 regions of the right hemisphere of the aparc parcellation showing the highest spectral PLI peak.** The regions shown (one per column) correspond to the 8 regions of the aparc parcellation with the highest peak PLI coupling (from left to right). The curves correspond to the average over subjects and voxels inside each region of PLI, coherence, power and flattened power respectively. The coupling spectra were calculated between the acoustic signal and brain oscillatory activity during stimulus encoding (active period). (A to H) show the Phase Lag Index (PLI) and (I to P) the Coherence coupling spectra, respectively, for natural normal rate (in blue), time-compressed (in green), and natural fast rate speech (in orange). To evaluate the possible effects of power modulations on the coupling, (Q) to (X) display the brain activity's power spectrum during the encoding period, and (Y) to (FF) show their flattened curve using FOOOF.

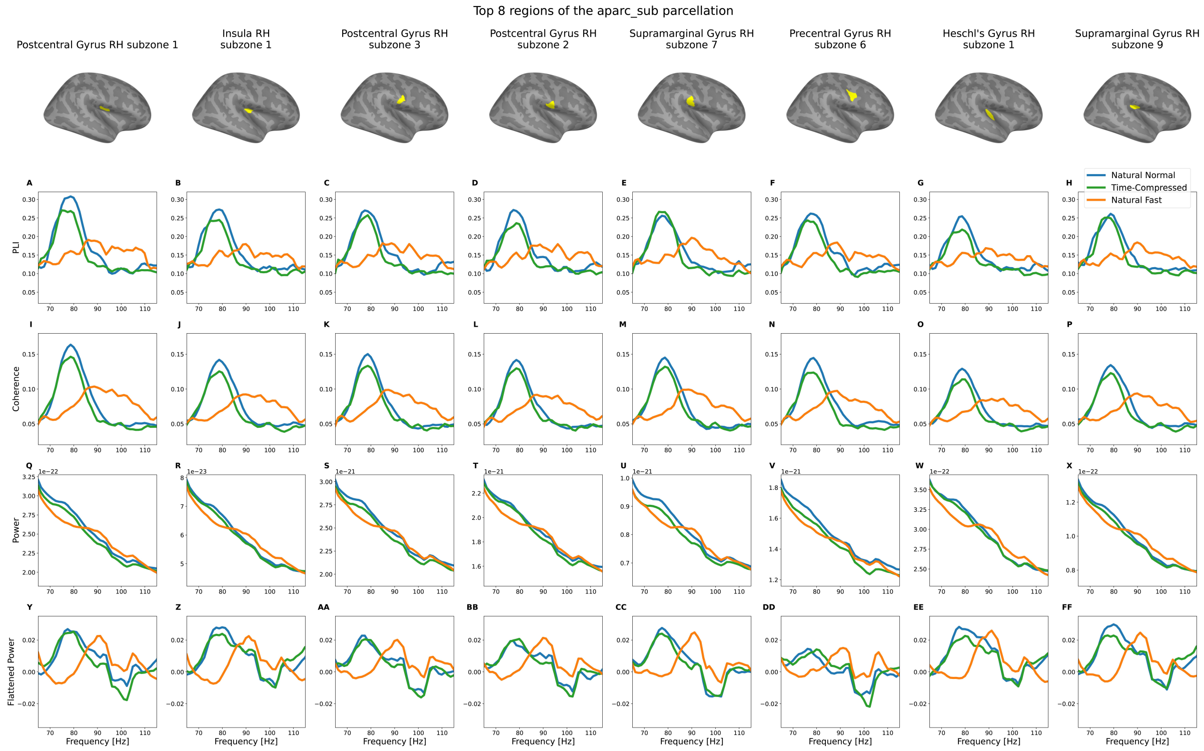


**Figure S3: Top 8 regions of the right hemisphere of the aparc_sub parcellation.** The regions shown (one in each column) correspond to the 8 regions of the aparc_sub parcellation with the highest peak PLI coupling (from left to right). The curves correspond to the average over subjects and voxels inside each region of PLI, coherence, power and flattened power respectively. The coupling spectra were calculated between the acoustic signal and brain oscillatory activity during stimulus encoding (active period). (A) to (H) show the Phase Lag Index (PLI) and (I) to (P) the Coherence coupling spectra, respectively, for natural normal rate (in blue), time-compressed (in green), and natural fast rate speech (in orange). To evaluate the possible effects of power modulations on the coupling, (Q) to (X) display the brain activity's power spectrum during the encoding period, and (Y) to (FF) show their flattened curve using FOOOF.

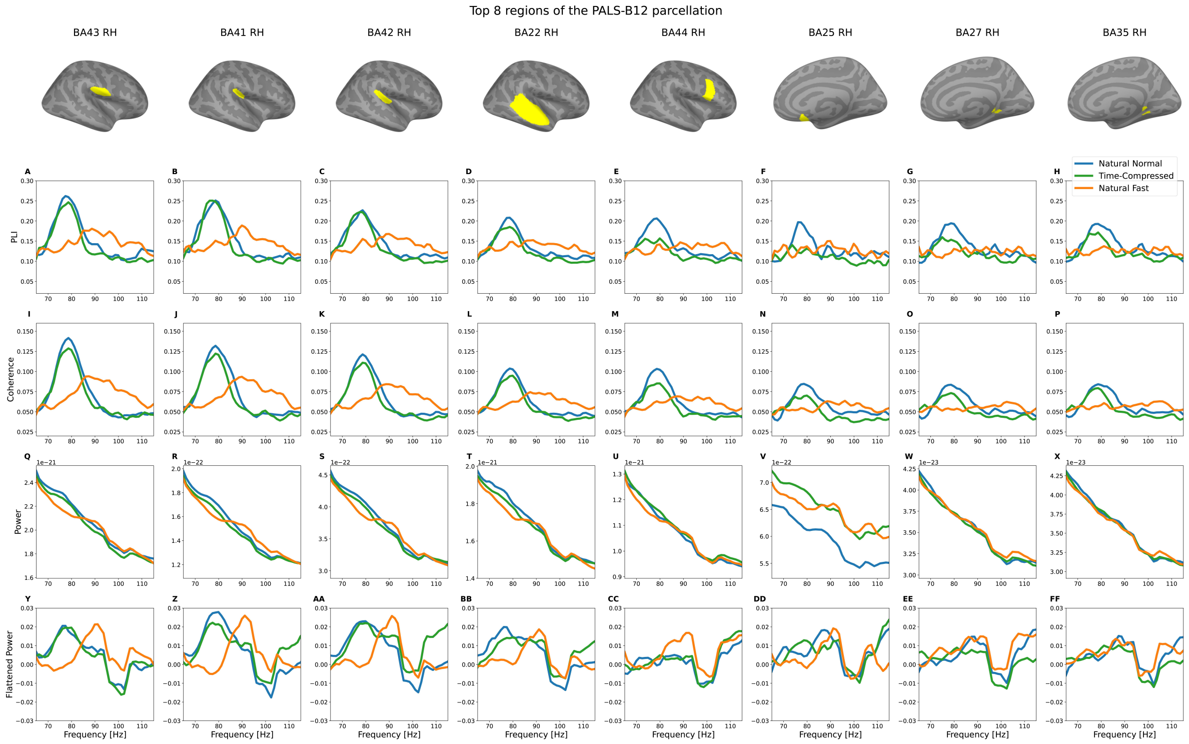


**Figure S4: Top 8 regions of the right hemisphere of the PALS-B12 parcellation.** The regions shown (one in each column) correspond to the 8 regions of the PALS-B12 parcellation with the highest peak PLI coupling (from left to right). The curves shown correspond to the average over subjects and voxels inside each region of PLI, coherence, power and flattened power respectively. The coupling spectra were calculated between the acoustic signal and brain oscillatory activity during stimulus encoding (active period). (A) to (H) show the Phase Lag Index (PLI) and (I) to (P) the Coherence coupling spectra, respectively, for natural normal rate (in blue), time-compressed (in green), and natural fast rate speech (in orange). To evaluate the possible effects of power modulations on the coupling, (Q) to (X) display the brain activity's power spectrum during the encoding period, and (Y) to (FF) show their flattened curve using FOOOF.

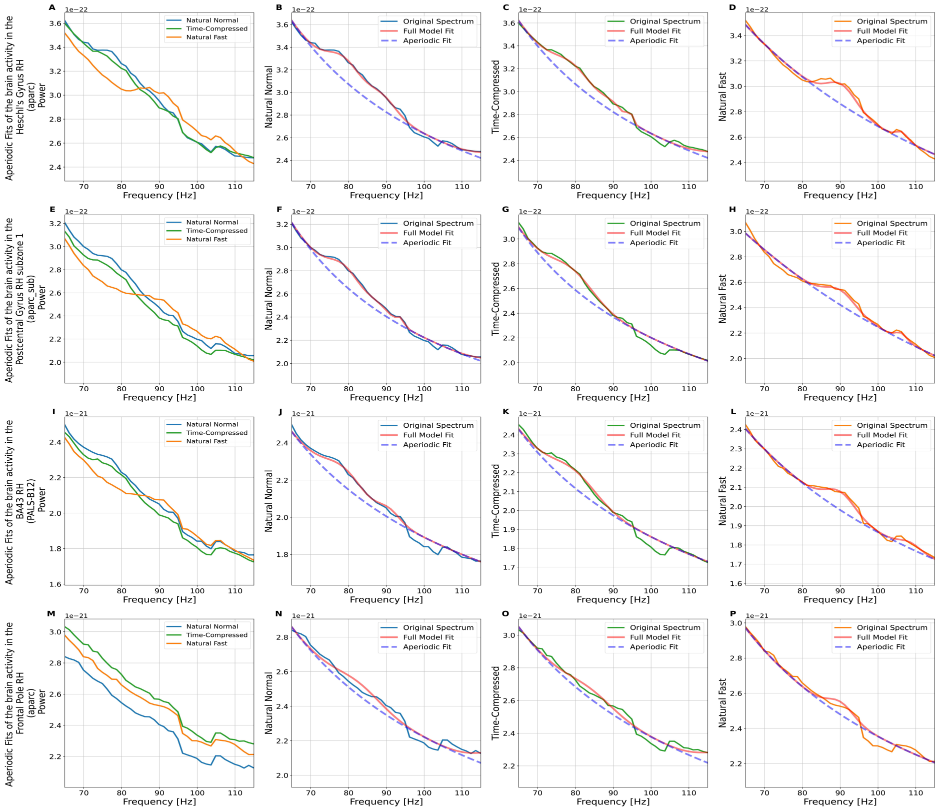


**Figure S5: Aperiodic fits of the mean brain activity power spectra in each region.** (A), (B), (C), (D) correspond to the right Heschl's gyrus (from aparc parcellation)**.** (A) Power spectra for the three speech conditions (same as Figure 3I; normal rate in blue, time-compressed in green and fast rate in orange). (B), (C) and (D) show the fitting using the FOOOF procedure (Donoghue et al., 2020) for normal rate, time-compressed and fast rate conditions, respectively. On each, the purple dashed line represents the aperiodic component, the red line the full model fit (the sum of the periodic and aperiodic components), and the third line shows the original spectra of the respective speech condition following the same color code as in (A). Similarly, the plots are repeated for the other regions. The first subzone of the right Postcentral Gyrus (from aparc_sub parcellation) corresponds to (E), (F), (G), (H). The right Brodmann Area 43 (from PALS-B12 parcellation) corresponds to (I), (J), (K), (L). The right Frontal Pole is used as a control region (from aparc parcellation); it corresponds to (M), (N), (O), (P).
